# *Ixodes ricinus* nymphs prefer blood with *Borrelia* infection and low corticosteroid concentration

**DOI:** 10.1101/2024.10.02.616250

**Authors:** Tosca Vanroy, Bram Catfolis, Elin Verbrugghe, Kris Verheyen, Luc Lens, Diederik Strubbe, Frank Pasmans, An Martel, Siska Croubels, Marc Cherlet, Lander Baeten

**Affiliations:** Forest & Nature Lab, Department of Environment, Faculty of Bioscience Engineering, Ghent University, B-9090 Melle-Gontrode, Belgium; Terrestrial Ecology Unit, Department of Biology, Faculty of Sciences, Ghent University, B-9000 Ghent, Belgium; Wildlife Health Ghent, Department of Pathobiology, Pharmacology and Zoological Medicine, Faculty of Veterinary Medicine, Ghent University, B-9820 Merelbeke, Belgium; Laboratory of Pharmacology and Toxicology, Department of Pathobiology, Pharmacology and Zoological Medicine, Faculty of Veterinary Medicine, Ghent University, B-9820 Merelbeke, Belgium

**Keywords:** *Borrelia burgdorferi*, host, glucocorticoids, preference experiment, *in vitro*

## Abstract

Ticks play a significant role in the transmission of various pathogens, impacting both human and animal health. Understanding the factors influencing tick feeding preferences is crucial for mitigating the risk of tick-borne diseases. This study investigates the blood preference of *Ixodes ricinus* nymphs, focusing on host species, stress hormone levels (glucocorticoids), and the presence of *Borrelia burgdorferi* bacteria. Three series of *in vitro* experiments were conducted. The setup involved placing individual blood drops (15 µl) on filter paper over a 37°C plate. Ticks were placed in the center, and their movements were tracked for two minutes to record preferences. The first experiment evaluated the tick’s preference for blood from different hosts (mouse, bird, sheep). The second experiment explored the role of stress hormones by assessing the ticks’ response to blood with varying levels of supplemented cortisol and corticosterone (0, 10, 100 and 1000 ng/ml). The third experiment examined the potential influence of *Borrelia* infection, combined with elevated stress hormone levels, on blood preference. Our results show that *I. ricinus* nymphs show a significant preference for blood without added glucocorticoids and, in mice, blood with *Borrelia* infection. No clear preference for a specific host species was observed. These findings provide valuable insights into the complex interactions between ticks, blood characteristics, and *Borrelia* infection status, highlighting the potential importance of host physiological state in tick host selection. Future studies should explore the mechanisms behind these preferences, which could lead to the development of new methods to control the spread of tick-borne diseases.

**HIGHLIGHTS:**

- *In vitro* experiments with nymphs can reveal insights into their blood preferences
- Ticks displayed no difference in preference towards bird, mouse, or sheep blood
- Ticks prefer blood with baseline stress levels over increased stress levels
- Mouse blood infected with *Borrelia burgdorferi* was preferred by ticks

## INTRODUCTION

Ticks (Acari: Ixodidae) are hematophagous ectoparasites with a significant impact on both human and animal health through their role as vectors of various pathogens. To complete their life cycle, ticks need to feed on the blood of vertebrates, and in doing so, can transmit a variety of pathogens, including bacteria (e.g., *Borrelia*, *Anaplasma*), viruses (e.g., tick-borne encephalitis), and parasites (e.g., *Babesia*, *Theileria*) (Baneth, 2014; Boulanger et al., 2019; Brites-Neto et al., 2015; Dantas-Torres et al., 2012). In recent decades, climate change and human-induced land use changes have led to an increase in the abundance and geographical expansion of ticks, particularly in the northern hemisphere (Bouchard et al., 2019; Diuk-Wasser et al., 2021; Sonenshine, 2018; Wikel, 2018). As a result, tick-borne diseases have become more prevalent and widespread, posing a growing public health problem (Rochlin & Toledo, 2020). For example, Lyme borreliosis, caused by *Borrelia burgdorfer*i sensu lato, is the most common tick-borne disease in Europe and the USA (Marques et al., 2021). In Western Europe, the incidence of Lyme borreliosis is increasing, particularly in the northern and central regions (Burn et al., 2023; Vandekerckhove et al., 2021). The disease is also a major health issue in North America, with thousands of new infections reported each year (Dumic and Severnini, 2018; Matuschka and Spielman, 1986; Murphree Bacon et al., 1991; Nelson et al., 2015; Schwartz et al. 2017). Therefore, understanding the factors that influence the feeding preferences of ticks is crucial for developing effective strategies to mitigate the risk of tick-borne diseases.

Unlike insects, ticks do not have antennae as a sense organ. Instead, they use the Haller’s organ on the tarsus of their first pair of legs to locate possible hosts and conspecifics (Carr and Salgado, 2019; Lees, 1948). The Haller’s organ has olfactory (pheromones, kairomones and CO_2_), mechanosensory, hygrosensory (humidity) and thermosensory (temperature) functions (Carr and Salgado, 2019). Questing behaviour is usually characterized by a tick positioned on vegetation with its front legs extended upwards. This is done not only to grasp a host, but also to pick up cues from hosts with their Haller’s organ. For long-range detection, ticks use infrared radiation, vibrations and host odors(Carr and Salgado, 2019; Mitchell et al., 2017; Wanzala et al., 2004). When a host is closer, ticks also rely on moisture and less volatile chemical cues (Bezerra-Santos et al., 2024; Crooks and Randolph, 2006). Most studies have focused on finding chemical compounds that either repel (Bissinger and Roe, 2010; Faraone et al., 2019; Grenacher et al., 2001; Romashchenko et al., 2012) or attract (Dallas and Foré, 2013; Romashchenko et al., 2012) ticks, to find a way to reduce contact with ticks and tick-borne pathogens. However, little is known about host-associated attractants. Ticks are known to respond positively to CO_2_ (Steullet and Guerin, 1992; Van Duijvendijk et al., 2017a) and kairomones (Leonovich, 2004; Osterkamp et al., 1999; Soares and Borges, 2012), both of which are used to detect the presence of a host. Some vector-borne pathogens can alter the odor of a host, making it more attractive to vectors, as described for *Plasmodium* infections and mosquitos (De Moraes et al., 2014). Van Duijvendijk et al. (2017b) found that *Ixodes ricinus* nymphs were more attracted to the odour of *Borrelia afzelii*-infected bank voles (*Myodes glareolus*), but it was not clear whether infection led to a change in odour or an increase in CO_2_ production in the host.

Changing environmental conditions affect not only the prevalence of ticks and the incidence of tick-borne diseases, but also the communities of their hosts. Habitat fragmentation and degradation, for example, have been shown to induce a stress response in free-ranging vertebrates (Wingfield, 2013). This response is characterized by an immediate increase in catecholamine levels, followed by elevated glucocorticoid levels, such as cortisol or corticosterone, depending on the vertebrate species (Karaer et al., 2023; Madliger and Love, 2014). While short-term elevations of these plasma glucocorticoids are considered adaptive, long-term or chronic elevations can negatively affect individual fitness (Bonier et al., 2009; Breuner et al., 2008; Vitousek et al., 2018). These negative effects include delaying or reducing reproduction, suppressing growth, causing oxidative stress or impairing the immune system, and can therefore make individuals more susceptible to diseases (Costantini et al., 2011; Homyack, 2010). The immunosuppressive effects of chronic stress have been repeatedly demonstrated in mice and birds, the two main hosts of the immature stages of *I. ricinus* (Domínguez-Gerpe and Rey-Méndez, 2001; Frick et al., 2009; Kiank et al., 2006; Nazar and Marin, 2011). However, an effective immune response is crucial for managing and minimizing the impact of tick infestations and fighting potential infections with tick-borne pathogens, especially since tick saliva contains a cocktail of bioactive molecules designed to modulate host immunity and suppress its inflammatory responses (Chmelař et al., 2016; de la Fuente et al., 2016; Wikel, 2013). If questing ticks can detect stress levels before attaching, they could potentially select hosts with impaired immune function, leading to higher feeding success and shorter feeding times. For instance, Gervasi et al. (2016) showed that zebra finches (*Taeniopygia guttata*) with experimentally elevated corticosterone levels were approximately twice as likely to be fed on by mosquitoes. However, this crucial mechanism in ticks remains poorly understood.

To fill the knowledge gap on host preference by ticks, we performed a successive series of *in vitro* experiments, inspired by the work of Žákovská et al. (2018) on tick preferences for human blood groups. In their approach, drops of each blood type (A, B, AB, or O) were applied to sterile filter paper in a Petri dish, followed by the observation of a tick’s movements. This *in vitro* experimental design offers several advantages: it is simple, time-efficient and eliminates the need for animals, thereby excluding host behaviour as a confounding factor. In a first experiment, we test *I. ricinus* preference for blood from different hosts (mice, birds, and sheep), both with and without the addition of very low concentrations of ethanol. Ethanol is an essential solvent for glucocorticoids prior to blood spiking and therefore needs an evaluation to rule out any potential repellent properties. To examine the role of stress hormones in host selection, in a second experiment, we explore the ticks’ response to blood containing different levels of administered glucocorticoids (cortisol and corticosterone), from 0 to 10, 100 and 1000 ng/ml. To investigate the potential role of blood pathogens in tick feeding choices, a third experiment was designed to assess tick preference for blood infected with *Borrelia burgdorferi* sl., taking into account both baseline and elevated stress levels in blood. To address for inherent affinity of tick nymphs towards specific host types, the second and third experiment were performed on both mouse and avian blood, two main host types for *I. ricinus* nymphs.

## MATERIALS & METHODS

### Ticks

*Ixodes ricinus* nymphs were purchased from Insect Services GmbH (Berlin, Germany). They were at least 10 months old at the time of the experiments. Prior to the experiment, the ticks were housed in 15 ml plastic tubes with humified paper and sealed with fine mesh. They were kept in an incubator at 23°C and relative humidity between 85% and 95%, with a 14:10h light: dark regime. Only nymphs showing a negative geotaxis in their container were used for the experiments.

### Chemicals

Cortisol and corticosterone (Sigma-Aldrich, respectively H4001-1G and 27840-100MG) stock solutions of 10 mg/ml, 1 mg/ml and 100 µg/ml were prepared in 100% ethanol, aliquoted and stored at - 20°C. To spike the blood samples, stock solutions were diluted 1:100 in demineralised water and then further diluted 1:100 in blood to obtain concentrations of 1000, 100 and 10 ng/ml. By doing so, the volume of ethanol was kept constant in the different concentration conditions and minimally diluted the blood. We selected these concentrations to represent a spectrum of physiological stress responses. The 10 ng/ml concentration represents a slight elevation from baseline levels, while 100 ng/ml mimics moderate to severe stress, as supported by studies on stress-induced glucocorticoid levels in rodents and birds (Gong et al., 2015; Hillebrecht et al., 2024; Kim et al., 2013; Lee et al., 2011; Liu et al., 2013; Post et al., 2003; Scanes, 2016; Taban et al., 2016). The 1000 ng/ml concentration, although rarely encountered in nature, was included as an experimentally high level. A solvent control was prepared by diluting ethanol 1:100 in water and consequently 1:100 in blood. To spike BSK(Barbour-Stoenner-Kelly)-H (Sigma-Aldrich) medium with 2000 ng/ml cortisol or corticosterone, the 10 mg/ml stock solutions were diluted 1:5000 in BSK-H medium. A solvent control was prepared by diluting ethanol 1:5000 in BSK-H medium.

### *Borrelia burgdorferi* cultivation, quantification and glucocorticoid metabolization

An aliquot of frozen *Borrelia burgdorferi* sensu stricto (DSM 4680) was thawed at room temperature, transferred into T25 flasks with a vented cap containing 5 ml of complete BSK-H medium and grown for 5 to 7 days in a microaerobic environment. To examine the possible preference of *I. ricinus* nymphs for blood spiked with *B. burgdorferi* whether or not in combination with stress hormones, a 5-day old *B. burgdorferi* culture was 1:2 diluted in (A) unsupplemented BSK-H medium, (B) BSK-H medium with ethanol (0.02%, solvent control), (C) BSK-H medium with 1000 ng/ml cortisol or (D) BSK-H medium with 1000 ng/ml corticosterone. These cultures were then incubated for 24 hours in a microaerobic environment. To assess potential differences in *Borrelia* abundance across the different conditions, a small aliquot (100 µl) from each culture was used for DNA extraction using PrepMan, followed by PAN-Borrelia real-time PCR (Cull et al., 2021). The PCR result show a minimal variation between the cultures as indicated by a mean LOG_10_(gene copies/ml) of 6.34 with a standard deviation of ± 0.3. The rest of the cultures were centrifuged for 10 min at 4000 x g, and the bacterial pellet was resuspended in (A) unsupplemented blood (from bird and mouse) or in blood (from bird and mouse) supplemented with (B) ethanol (0.02%, solvent control), (C) 1000 ng/ml cortisol or (D) 1000 ng/ml corticosterone. Whole blood without *Borrelia* was included as a control. After a 1-hour incubation period at 37°C, the preference of *I. ricinus* nymphs for these five conditions was tested (see Blood preparation). To evaluate potential glucocorticoid metabolism by *Borrelia* in bird and mouse blood, a small aliquot of the 1-hour incubated *Borrelia*-supplemented blood was analyzed for corticosterone and cortisol concentrations using an ultra-high-performance liquid chromatography-tandem mass spectrometry method (UPLC-MS/MS) (Supplementary Materials S1). Additionally, baseline corticosterone and cortisol concentrations were measured in unsupplemented whole blood from bird, mouse, and sheep for comparison. These tests were run in duplicate. No significant metabolization of glucocorticoids by *B. burgdorferi* was detected (Table S2).

### Blood preparation

Chicken blood (*Gallus gallus domesticus*) was purchased from Poulpharm (Izegem, Belgium) and blood from outbred mice (*Mus musculus*) was purchased from Janvier Labs (Le Genest-Saint-Isle, France). Both blood samples were collected in lithium heparin-coated tubes to prevent clotting. Defibrinated sheep blood (*Ovis orientalis aries*) was acquired from BioTrading (Keerbergen, Belgium). Both anticoagulation methods have been shown not to impact tick feeding behaviour (Asri et al., 2023; Romano et al., 2018; Voigt et al., 1993; Waladde et al., 1993). All blood samples were aliquoted and stored at -20°C. Defrosted blood has been used successfully for feeding adults and nymphs of different species and has been show to obtain similar results compared to fresh blood (González et al., 2017, 2021; Krull et al., 2017).

For the first host-preference experiment, six distinct blood conditions were prepared. For each host type (mouse, bird, sheep), two blood samples were arranged: unsupplemented whole blood and a solvent control. In a second experiment, the possible preference for stress hormone-spiked blood was tested. This was both performed in mouse and chicken blood. Although blood was collected in lithium heparin tubes, some clotting was observed in chicken blood upon thawing. To mitigate this, we added lithium heparin (2 µl per 1 ml of blood), a method previously demonstrated to not adversely affect artificial tick feeding (Asri et al., 2023; Liu et al., 2014; Romano et al., 2018; Voigt et al., 1993; Waladde et al., 1993). Per host species, five distinct blood conditions were prepared: blood containing 10, 100 and 1000 ng/ml stress hormone, a solvent control and unsupplemented whole blood. This was tested both for cortisol and corticosterone. In a third experiment, we examined whether nymphs show a preference for blood spiked with *B. burgdorferi* and/or stress hormones. We selected the highest stress hormone condition (1000 ng/ml) in this trial, resulting in 5 conditions: unsupplemented whole blood, or blood supplemented with *Borrelia*, *Borrelia* + 1000 ng/ml cortisol, *Borrelia* + 1000 ng/ml corticosterone, *Borrelia* + ethanol (solvent control). This was both performed in mouse and chicken blood.

### Experimental setup

The experiments were performed between July and November 2023. The experimental setup was based on the method described by Žákovská et al. (2018) with some adaptations and clarifications (Figure 1). The lid of a sterile Petri dish was placed upside down on a hot plate at 40°C, simulating an animal environment. On top of this lid, a sterile round filter paper was placed, and individual drops of blood (15 µl) for each condition were applied by a first researcher. To prevent attractants, such as sweat, adhering to the filter paper, both researchers wore gloves during the entire experiment. The drops were equally spaced and positioned 3 cm away from the middle. The drops were applied at the same positions within the Petri dish each time, which were letter-coded (e.g. A-E in a five-level treatment experiment) and treatments were randomly applied across replicates, to avoid a confound between position and treatment location. For each replicate, the position of each treatment was recoded. A second researcher placed one active tick in the middle of the filter paper and observed its movements for 2 minutes. The initial preference at time 0 (t_0_) was determined as the first direction in which the tick moved. To further assess the tick’s preference, the circle was divided into 5 or 6 equal parts, and the tick’s location was noted based on the section of the circle it occupied at minute one (t_1_) and two (t_2_). If the tick did not show a clear preference for any location at a certain time, it was marked as “No Choice” (NA) in the dataset. Subsequently, the tick was euthanized by immersion in ethanol. For each experiment, this protocol was repeated 100 times.

**Figure 1:**
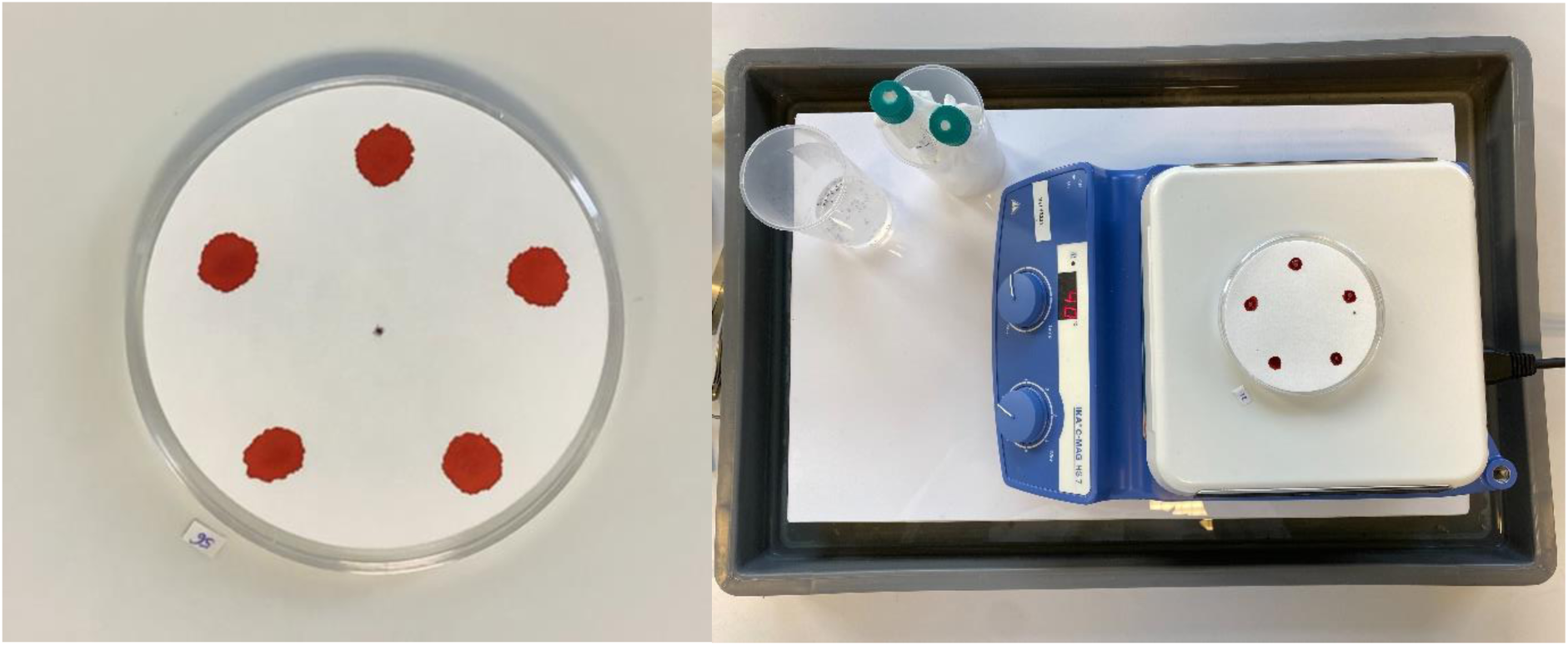
View of the setup of the experiments. Left: The Petri dish with five blood drops and a nymph *Ixodes ricinus* in the middle. Right: the full setup of the experiment with the Petri dish on the hot plate.

### Statistical analysis

A multinomial logistic regression was performed for each of the seven experiments using the brms-package (Burner, 2017) in R 4.3.1 (Team, 2023). The treatment of the selected blood drop at each timepoint was used as response variable. We used a multilevel (mixed) model to account for the spatio-temporal dependencies in the data. The multiple observations of each replicate dish (i.e. three timepoints) was dealt with by including the ID of each dish as a group-level (random) effect. A multinomial logistic regression with the chosen position as the response variable was constructed to investigate if there was a preference for a particular position within the Petri dish. We found that in all seven experiments at least one location was chosen more or less often than expected by chance, indicating directionality independent of the treatments (Figure S1-S7). The preference for certain locations differed between the different experiments, which could be due to different conditions in the lab on different days, such as air flow. To correct for this preference for certain locations, we added the position of the chosen drop as an additional crossed group-level (random) effect.

## RESULTS

### Experiment 1: Impact of ethanol on tick blood selection across three host species

When comparing the preference for blood between the three host species (sheep, mouse and bird) with and without ethanol added (solvent control), we found that sheep blood with ethanol was chosen slightly less than expected (at the 68% Credible Interval), whereas sheep blood without ethanol was slightly more preferred than expected (at the 68% CI) (Figure 2A). When considering only the three host species and combining the options with and without ethanol, no statistically clear difference in preference was observed. Nonetheless, there appears to be a trend towards preference for mammalian blood over avian blood (Figure 2B). Similarly, when analyzing the differences in choice between the conditions with and without ethanol, a preference for blood without ethanol may be noted. However, the difference between these two options was not statistically clear (Figure 2C).

**Figure 2:**
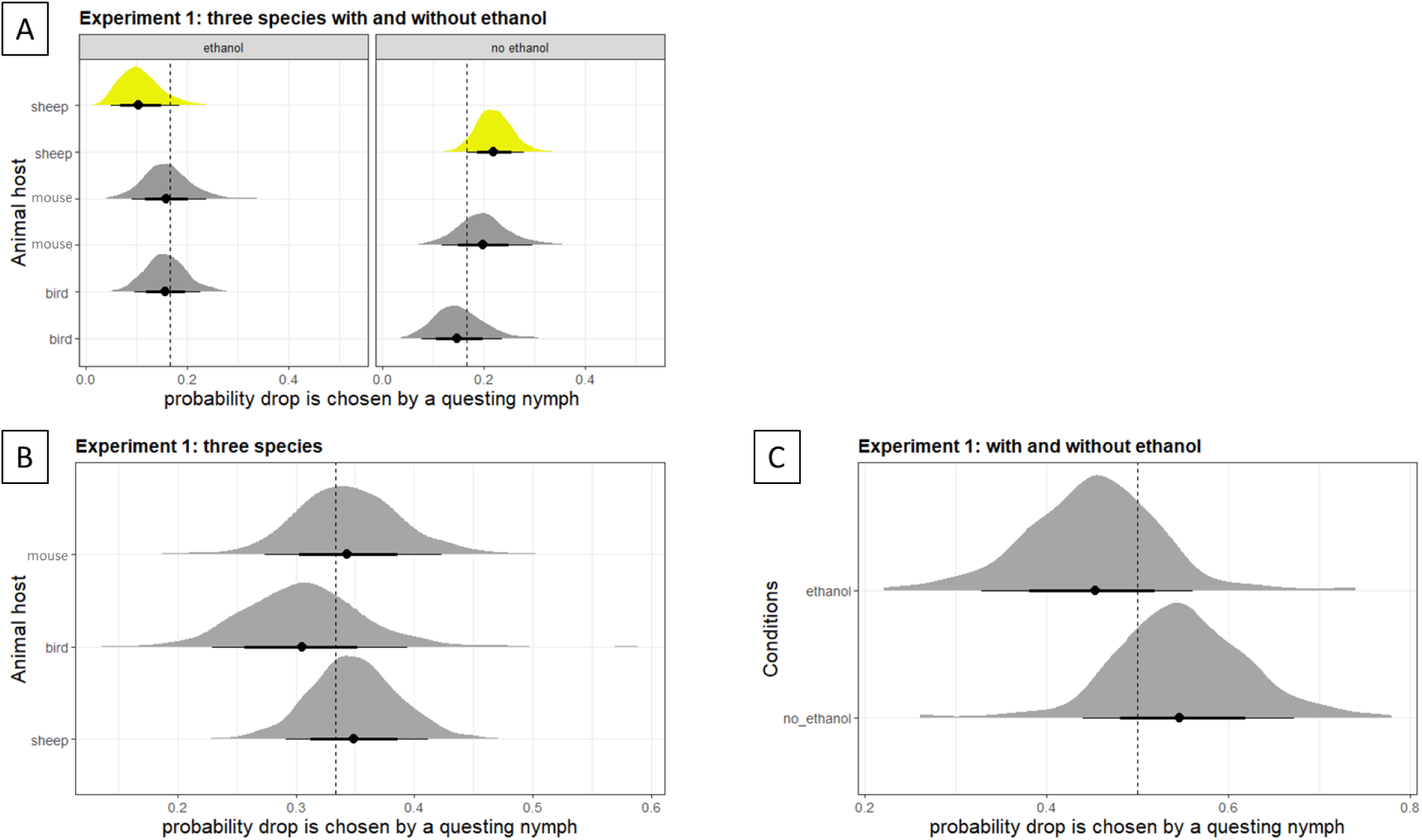
Tick preference for sheep, mouse or bird blood, with or without ethanol (experiment 1). A) The difference in preference between all six options, B) the probability that one of the three host species was chosen, independent of the presence or absence of ethanol (i.e. pooling with and without ethanol treatments), and C) the probability that a blood drop with or without ethanol was chosen, independent of the species (i.e. pooling across species). Graphs show posterior distributions of estimated probabilities of a drop being chosen. The dot represents the median of the distribution with 90% (thin lines) and 68% (thick lines) credible intervals (CI). Dashed line is the expected probability when there is no preference (i.e., 1 / #treatment levels). Yellow distributions are different from this expectation at the 68% CI. See Table S3 for median (+/- 68% quantile) values.

### Experiment 2: Preference patterns in cortisol and corticosterone supplemented mouse and bird blood

For mouse blood spiked with cortisol, we found that the solvent control and the 1000 ng/ml concentration were selected less frequently (at 68% CI) than expected. Conversely, unsupplemented whole blood was selected more frequently than the average, also within the 68% CI (Figure 3A). When corticosterone was added to mouse blood, it was found that the absence of administered corticosterone (whole blood & solvent control) was favored more frequently than expected, while concentrations of 100 ng/ml and 1000 ng/ml were selected less often (Figure 2B). These findings suggest a preference for lower levels of corticosterone in mouse blood. In the experiments conducted with bird blood, a lower preference for 100 ng/ml cortisol was observed (Figure 3C). No discernible difference in preference for blood supplemented with corticosterone was detected (Figure 3D).

**Figure 3:**
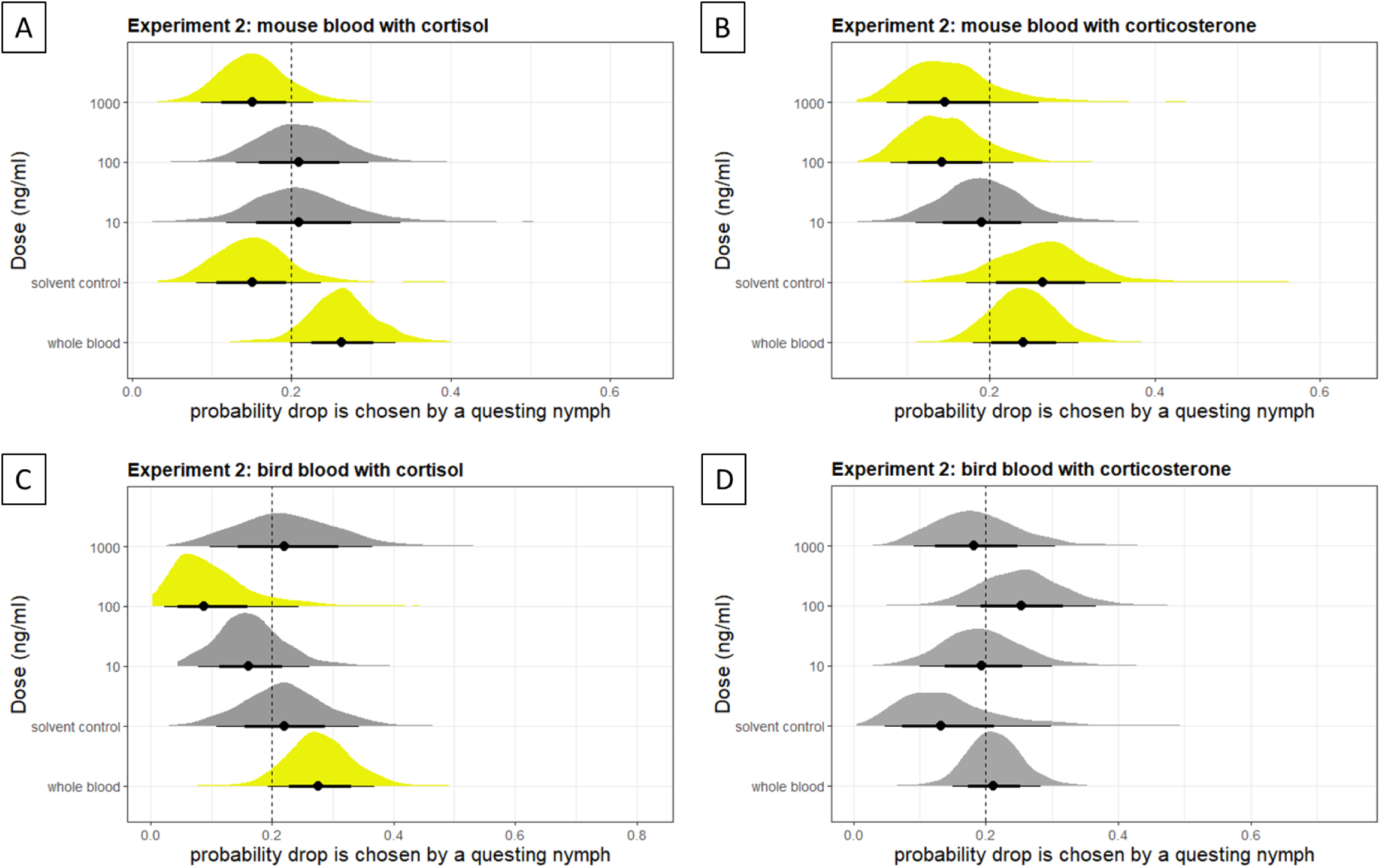
Tick preference for increasing glucocorticoid levels (whole blood, solvent control, 10, 100 and 1000 ng/ml) in mouse and bird blood (experiment 2). A) mouse blood with cortisol, B) mouse blood with corticosterone, C) bird blood with cortisol and D) bird blood with corticosterone. Graphs show posterior distributions of estimated probabilities of a drop being chosen. The dot represents the median of the distribution with 90% (thin lines) and 68% (thick lines) credible intervals (CI). Dashed line is the expected probability when there is no preference (i.e., 1 / #treatment levels). Yellow distributions are different from this expectation at the 68% CI. See Table S3 for median (+/- 68% quantile) values.

### Experiment 3: Preference patterns in *Borrelia*-infected mouse and bird blood with baseline or elevated glucocorticoid levels

In the experiment involving mouse blood, we found a higher preference (at 68% CI) for blood with *Borrelia* alone and a lower preference for blood containing *Borrelia* combined with 1000 ng/ml corticosterone (Figure 4A). For bird blood, no significant difference in preference among all five options was discerned (Figure 4B).

**Figure 4:**
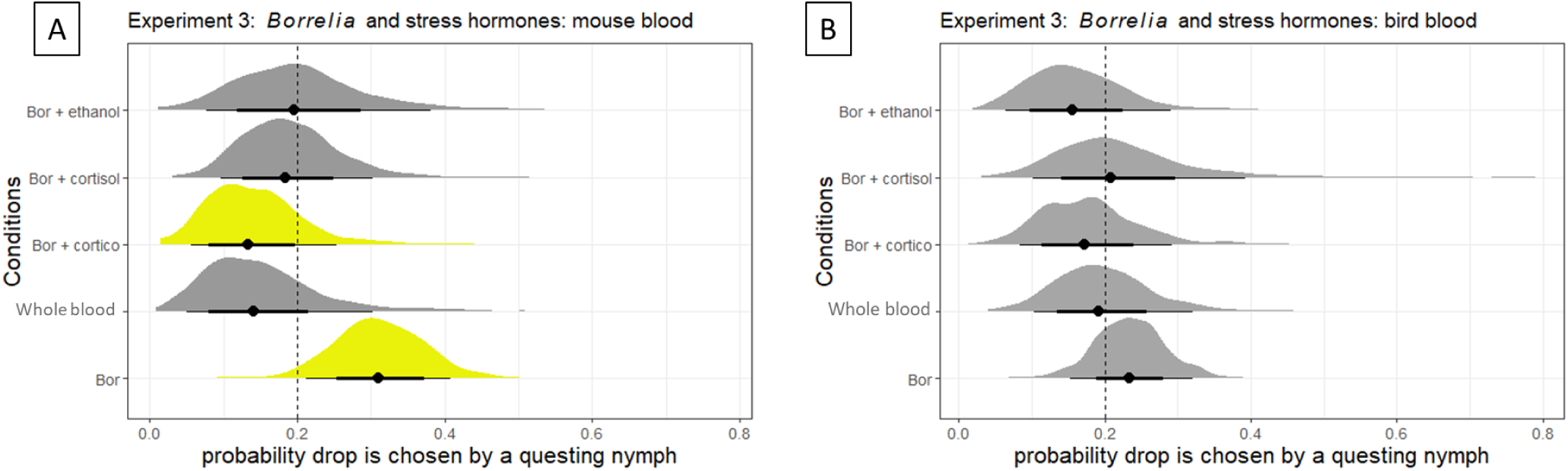
Tick preference for *Borrelia* (Bor) with or without high levels (1000 ng/ml) of stress hormones (cortisol and corticosterone (cortico)) (experiment 3) in A) mouse blood and B) bird blood. Graphs show posterior distributions of estimated probabilities of a drop being chosen. The dot represents the median of the distribution with 90% (thin lines) and 68% (thick lines) credible intervals (CI). Dashed line is the expected probability when there is no preference (i.e., 1 / #treatment levels). Yellow distributions are different from this expectation at the 68% CI. See Table S3 for median (+/- 68% quantile) values.

## DISCUSSION

Our study investigated how host species, blood glucocorticoid levels and *B. burgdorferi* infection influence the selection behaviour of questing *I. ricinus* nymphs. To our knowledge, the scientific literature has not yet explored whether ticks exhibit differential preferences based on these blood characteristics (Defolie et al., 2020). This experiment thus aimed to fill this gap in our understanding of tick host selection, and could have significant implications for understanding tick-borne disease transmission. In general, our results points towards a preference of ticks for blood with baseline glucocorticoid levels (most profound in mice) and *Borrelia-*infected blood (only in mice). No general significant effect was observed for very low ethanol concentrations in blood. Here, we further discuss the influences of host species, stress levels, and *Borrelia* infection on tick preference.

### No effect of host species on blood selection

Our first experiment found no significant preference among *I. ricinus* ticks for the three host species tested (chicken, mouse, and sheep). This supports the established view of this tick species as a generalist vector, capable of utilizing a large variety of hosts (Gray et al., 2021). Accordingly, in an artificial feeding experiment, no significant differences were found in the proportion and weight of engorged ticks, nor in the duration of feeding, between chicken and sheep blood (Liu et al., 2014). However, some studies suggest potential host preferences within tick populations. Kempf et al. (2011) identified genetic differences between tick populations feeding on birds and rodents, hinting at possible adaptations to specific hosts. Additionally, Heylen et al. (2010) reported that the mean weight of *I. ricinus* nymphs fed on mice was 3.7 mg, whereas the mean weight of bird-fed nymphs has been reported as 4.2 mg. Further research through methodical comparative studies on the long-term performance (engorgement, development, egg laying) of *I. ricinus* ticks feeding on different host species is required.

While very low concentrations of ethanol did seem to slightly impact *I. ricinus* nymphs’ selection of sheep blood, it did not significantly affect their overall blood preference. It is important to note that ethanol was only used at very low levels (0.02%) to dissolve the glucocorticoids, which are poorly soluble in water. Previous experiments have tested the toxicity and repellence of ethanol towards ticks, but only at high concentrations of ethanol (Gonçalves et al., 2007; Romashchenko et al., 2012). Therefore, the low levels used in our experiment are unlikely to have a significant impact on tick preference.

### Preference for baseline glucocorticoid levels

Contrary to our expectations, our results indicate that ticks, specifically *I. ricinus* nymphs, may be more likely to select hosts with lower, baseline concentrations of glucocorticoids in their blood and, as such, represents the first investigation into tick host selection based on varying glucocorticoid levels in blood.

In mice, this trend was more pronounced for blood spiked with corticosterone compared to cortisol. While glucocorticoids themselves are not known to directly affect arthropods, this difference could be attributed to the different roles of these hormones in stress regulation within the host. Although mouse serum cortisol and corticosterone dynamics are closely correlated across various physiological or stressful situations, research suggests they have distinct functions (Gong et al., 2015). Cortisol acts as a faster responder during acute stress situations (Gong et al., 2015), while corticosterone appears to be more closely associated with the body’s adaptation to chronic stress, influencing factors like body mass and food intake (Jeong et al., 2013). By preferentially selecting hosts with lower corticosterone levels, ticks might be indirectly selecting for healthier individuals experiencing less chronic stress. This could benefit the tick’s own survival and reproductive success in several ways. Healthier hosts with lower stress might have higher nutritional value for the tick, leading to better growth and development (Bize et al., 2008; Christe et al., 2003; Maaz et al., 2018). A healthier host with a stronger immune response might be able to carry the tick for a longer period, allowing for a more complete feeding cycle, which could ultimately lead to a higher chance of successful reproduction for the tick (Heylen and Matthysen, 2011; Rynkiewicz et al., 2013; Václav et al., 2008).

Surprisingly, in birds, we observed a clear preference pattern only in cortisol-treated blood, while no distinct pattern emerged in corticosterone-treated samples. This is contradictory to our expectations given the fact that corticosterone is considered the dominant glucocorticoid in birds, as well as in rodents (Botía et al., 2023). A possible explanation might be a high baseline level of corticosterone in bird blood. However, our results show a low baseline corticosterone level in bird blood compared to mouse blood (Table S2). Further research is needed to understand this discrepancy between bird and rodent responses.

### Preference for *Borrelia*-infected blood in mice

*I. ricinus* nymphs seem to have a preference for blood infected with *B. burgdorferi*. Similar results were found in an olfactory experiment where *I. ricinus* nymphs preferred the odor of bank voles infected with *B. afzelii* over non-infected individuals (Van Duijvendijk et al., 2017b). The preference for an infected individual might be due to a modification of the host’s odor or increased CO_2_ production. However, since our study only used blood, these factors do not explain our findings. There is substantial evidence that vectors prefer infected hosts, a phenomenon observed across various host species, particularly in mosquitoes (Cozzarolo et al., 2020; Lacroix et al., 2005; Zhang et al., 2022). Mathematical models show that stable infections are easier to maintain when vectors have a preference for infected hosts, than when they randomly select hosts (Kingsolver, 1987). However, vectors cannot afford to be too selective, especially when host densities are low.

Adding high concentrations of glucocorticoids can influence the preference for *Borrelia*-infected blood, at least for high levels of corticosterone in mouse blood. This is in line with the findings of our second experiment, where we found that nymphs prefer baseline over increased stress levels.

### Limits and recommendations for further research

While the simplicity of the experimental design, as described by Žákovská et al. (2018), offered advantages in terms of ease of implementation and replicability, it also introduced certain limitations that warrant further investigation.

In this experiment, ‘preference’ is defined by the tick’s location at time t_x_, acknowledging the constraints posed by ticks’ inability to feed directly from blood drops. A next step could involve employing artificial membrane-based feeding systems to confirm whether ticks truly feed based on blood type (Asri et al., 2023; Böhme et al., 2018; Kröber and Guerin, 2007).

Ticks use chemical and thermal signals emitted by humans and other vertebrates to locate suitable hosts for a blood meal. Despite maintaining a distance of at least 1.5 meters, CO_2_ exhaled by observers might have influenced tick behaviour. We found that some positions on the Petri dish were selected more or less often than expected by chance, meaning that some locations were preferred due to unknown and uncontrolled environmental factors. This preference for certain positions on the Petri dish varied between the different experiments, so we corrected for this in the statistical analysis by including the location of the drop as a random factor. Species like *I. scapularis* (Städele, 2024) and *Amblyomma variegatum* (Steullet and Guerin, 1992) are known to be highly sensitive to CO_2_ as a long-range host attractant, even at low concentrations. Notably, research suggests that the detection of CO_2_ in *I. ricinus* nymphs acts as an activator of host-seeking behaviour. Host odour, potentially emanating from observers, then serves as the attractant, guiding the nymphs to select suitable questing locations and attachment sites on the host (Van Duijvendijk et al., 2017a). Future studies could thus benefit from employing a closed system or utilizing alternative methods to minimize tick exposure to kairomones.

Using their Haller’s organ, questing ticks also react to thermal stimuli, namely convective heat (i.e., hot air currents) and radiant heat (i.e., IR electromagnetic waves), emitted by an approaching animal host. This ability has been documented in several tick species, such as *Amblyomma americanum*, *Dermacentor variabilis* and *I. scapularis*, with some even detecting human hosts from several meters away (Carr and Salgado, 2019; Mitchell et al., 2017; Otálora-Luna et al., 2022). The sensitivity of *I. ricinus* to thermal stimuli remains untested, so further investigation is necessary to determine its influence on their behaviour in our experiment. However, the heat generated by the hot plate, mimicking the animal environment, likely exceeded the radiant heat emitted by the observers (Hardy, 1939). Therefore, we argue that the observed tick behaviour might be primarily driven by the stronger thermal stimulus from the hot plate rather than human presence.

This study provides novel insights into the influence of blood characteristics on *I. ricinus* nymph selection behaviour. However, several avenues warrant further exploration to broaden our understanding of this complex process. While this study examined blood from sheep, lab mice, and chickens, ticks encounter a wider variety of potential mammalian and avian hosts in nature. Investigating tick selection across ecologically relevant species, particularly deer and forest-dwelling rodents and birds, would provide a clearer understanding of their host selection and its potential impact on tick population dynamics.

This study also identified correlations between different stress hormone concentrations and tick selection. Future research should delve deeper into the underlying physiological and behavioural mechanisms driving these preferences. Do these glucocorticoids influence tick sensory perception, attachment success, or blood meal suitability? *In vivo* and field-based experiments coupled with physiological analyses are crucial to elucidate these causal links.

We further revealed a preference for *Borrelia*-infected blood in mice. Understanding the ecological and evolutionary significance of this finding is critical. Do these preferences translate into enhanced *Borrelia* transmission rates in certain host species? Longitudinal studies incorporating entire tick life cycles and transmission dynamics are needed to address this question.

By applying a simple and reproducible design, our study fills significant gaps in the literature and sets the stage for more detailed investigations into the multifaceted factors influencing *I. ricinus* host selection. Future research can build on our findings to develop targeted strategies for managing tick populations and reducing the incidence of tick-borne diseases.

## Authors’ contributions

**Tosca Vanroy**: Conceptualization, Methodology, Validation, Visualization, Formal analysis, Investigation, Data curation, Writing – original draft. **Bram Catfolis**: Conceptualization, Methodology, Validation, Visualization, Formal analysis, Investigation, Data curation, Writing – original draft. **Elin Verbrugghe**: Conceptualization, Methodology, Validation, Formal analysis, Investigation, Resources, Writing – review & editing. **Kris Verheyen**: Conceptualization, Methodology, Validation, Funding Acquisition, Writing – review & editing. **Luc Lens**: Conceptualization, Methodology, Validation, Funding Acquisition, Writing – review & editing. **Diederik Strubbe**: Conceptualization, Methodology, Validation, Investigation, Funding Acquisition, Writing – review & editing, Visualization. **Frank Pasmans**: Conceptualization, Methodology, Validation, Funding Acquisition, Writing review & editing, Supervision. **An Martel**: Conceptualization, Methodology, Validation, Funding Acquisition, Writing – review & editing, Supervision. **Siska Croubels**: Methodology, Validation, Investigation, Resources, Writing – review & editing. **Marc Cherlet**: Methodology, Validation, Investigation, Resources, Writing – review & editing. **Lander Baeten**: Conceptualization, Methodology, Validation, Formal analysis, Funding Acquisition, Writing – review & editing, Supervision.

## Data availability

The datasets generated and analyzed during the current study will be made available after publication.

## Acknowledgements

We thank Sofie De Bruyckere and Sarah Van Praet for their excellent technical input, including finetuning the protocol, housing the ticks, and growing and sequencing *Borrelia*. We are grateful to Ard Nijhof for providing valuable review comments. Corticosterone and cortisol were determined using an UPLC-MS/MS instrument part of the Ghent University MSsmall expertise centre for mass spectrometry analysis of small organic molecules, and partly funded by a Research Foundation Flanders (FWO) grant FWO.HMZ.2022.0008.01.

## SUPPLEMENTARY MATERIAL

**S1: Analysis of corticosterone and cortisol in whole blood**

### Chemicals and reagents

Corticosterone and cortisol analytical standards were respectively from Sigma-Aldrich and Supelco, purchased at Merck (Overijse, Belgium). Corticosterone-d8 and cortisol-d4 analytical standards (used as internal standard (IS)) were both from TRC (Toronto Research Chemicals, Toronto, Canada). Individual stock solutions of all components were prepared at 1 mg/ml in methanol and stored at ≤ -15 °C. For sample analysis, working solutions were prepared by appropriate dilution of the stock solutions in methanol. These working solutions were stored at 2 – 8 °C. A mixed working solution of corticosterone-d8 and cortisol-d4 both at a 100 ng/ml level was used in sample preparation. Mixed working solutions of corticosterone and cortisol in the concentration range 5 – 2500 ng/ml were used for spiking calibrator and quality control (QC) samples.

Methanol, used in stock- and working solution preparation and sample extraction, was of HPLC grade (Fisher Scientific, Filter Service, Eupen, Belgium). Water of Milli-Q grade was used for the preparation of the aqueous mobile phase component and for sample preparation, which was produced in-house by a water purifying system Milli-Q-SP (Sigma-Aldrich). Acetonitrile as the organic mobile phase component and formic acid as a mobile phase additive, were of ULC/MS grade (Biosolve, Valkenswaard, the Netherlands). Ethyl acetate involved in sample extraction was of *pro analysi* grade (Emsure_®_) (Merck).

### Sample preparation

To a 250 µl whole blood sample transferred quantitatively in a 15 ml plastic centrifuge tube, 25 µl of a 100 ng/ml corticosterone-d8 + cortisol-d4 mixed IS working solution was added, as well as 25 µl of methanol, followed by vortex mixing. The latter volume was replaced by 25 µl of a working solution of corticosterone and cortisol at varying levels for calibrator and QC sample preparation. For the preparation of matrix-matched calibrator and QC samples, and of blank samples as well, in-house blank chicken or mouse whole blood was used. Blank samples were not spiked with IS nor analyte, but instead 50 µl of methanol was added to these samples, before they were subjected to the sample preparation procedure.

Extraction consisted of a liquid-liquid extraction with ethyl acetate. Therefore, 3 ml of ethyl acetate was added to the 15 ml centrifuge tube, which was then put for 20 min at 33 rpm on a roller mixer SRT2 (Stuart Scientific, Novolab, Geraardsbergen, Belgium). Thereafter, the sample was centrifuged (3000 rpm, 10 min., 4 °C). The supernatant was transferred to a 15 ml glass tube, and dried under a gentle stream of nitrogen at 40 °C in a Pierce Reacti-Therm III™ Heating Module and Reacti-Vap™ III Module (Rockford, USA). The extract was redissolved first in 125 µl of methanol, followed by thorough vortex mixing. A 125 µl portion of water was further added, followed by a new vortex mixing step. The final extract was passed over a Nylon filter (0.22 µm, 13 mm) (Merck), and collected in a plastic conical vial, followed by injection of a 5 µl sample aliquot onto the UPLC-MS/MS system.

### UPLC-MS/MS analysis

The UPLC instrument consisted of an Acquity H-Class Quaternary Solvent Manager and an Acquity FTN Sample Manager (Waters, Milford, USA). Chromatographic separation was achieved on an Acquity UPLC_®_ BEH C18 column (1.7 µm, 50 x 2.1 mm i.d.), in combination with a precolumn of the same type (VanGuard_TM_, 5 mm x 2.1 mm i.d.), both from Waters. A gradient elution was performed with a mobile phase of 0.1% (v/v) formic acid in water (A) and 0.1% (v/v) formic acid in acetonitrile (B), at a flow rate of 0.3 ml/min, i.e. 0 min., 80% A/20% B; 0 – 3.0 min., to 20% A/80% B; 3.0 – 4.4 min., 20% A/80% B; 4.4 – 4.5 min., to 80% A/20% B; 4.5 – 6.0 min., 80% A/20%. The column was maintained at a temperature of 40 °C, while the autosampler was set at 10 °C. The total chromatographic run time was 6.0 min. The UPLC effluent was sent from 1.8 min to 3.5 min by use of a divert valve to a Xevo TQ-S_®_ triple quadrupole mass spectrometer from Waters, equipped with an ESI ion source operating in the positive ionization mode. The UPLC-MS/MS analysis was run under control of MassLynx software (v4.1), which was used for subsequent data processing as well. Operating conditions for the ESI source used in the positive ionization mode were optimized by direct infusion of all individual components in combination with the mobile phase at 50% A/50% B delivered at a flow rate of 0.3 ml/min. The following tune parameters were used for detection of all components: capillary voltage, 3 kV; cone voltage, 45 V; source offset, 50 V; source temperature, 150 °C; desolvation temperature, 500 °C; desolvation gas flow, 800 l/h; cone gas flow, 150 l/h; collision gas flow, 0.2 ml/min; ion energy 1, 0.5; ion energy 2, 1.5; LM 1 and LM 2 resolution, 2.8; HM 1 and HM 2 resolution, 14.8. Components were detected in MS/MS mode using component specific MRM (Multiple Reaction Monitoring) transitions, see Table S1. For corticosterone and cortisol, two specific product ions were monitored, a quantifier ion used for quantification purposes, and a qualifier ion, used for identification purposes, based on the ion ratio of both product ions. Since both corticosterone and cortisol are endogenous components, obtained quantification results were corrected for the endogenous levels present in the blank whole blood used for the construction of the calibration curve (response (peak area analyte/peak area IS) against the analyte concentration). The endogenous concentration (C_endo_) was obtained as the negative intersection with the x-axis of the calibration curve, calculated as C_endo_ = b/a, with a and b being respectively the slope and the intercept of the calibration curve.

**Table S1.** MS/MS detection parameters of all components.

| Component | Precursor ion ($m/z$) | Product ions ($m/z$) | CE (eV) |
| --- | --- | --- | --- |
| Corticosterone | 347.23 | Quantifier: 121.15 | 23 |
|  |  | Qualifier: 97.06 | 25 |
| Corticosterone-d8 (IS) | 355.32 | 125.2 | 23 |
| Cortisol | 363.26 | Quantifier: 121.00 | 25 |
|  |  | Qualifier: 327.12 | 15 |
| Cortisol-d4 (IS) | 367.29 | 121.01 | 25 |

*CE = Collision Energy (eV) used in fragmentation; $m/z$ = mass-to-charge ratio*

### Method validation

The UPLC-MS/MS analysis method for the analysis of corticosterone and cortisol in whole blood was validated using EC (2002/657/EC) and VICH (VICH GL49) guidelines.

For chicken whole blood, containing low endogenous levels of both corticosterone and cortisol, linearity in the range of 0.5 – 20 ng/ml was evaluated, with 6 calibrator levels being included, i.e. 0.5, 1, 2, 5, 10, and 20 ng/ml. Accuracy and precision were evaluated at a low and high concentration level, resp. 0.5 and 10 ng/ml, each on n= 6 replicates. The lowest level evaluated for accuracy and precision, i.e. 0.5 ng/ml, was established as the lower limit of quantification (LOQ) of the analysis method for both corticosterone and cortisol.

Mouse whole blood contains also a low endogenous level of cortisol, but on the other hand a high endogenous level of corticosterone. Therefore, for cortisol, the same concentration levels as above were involved at method validation. For this reason, the same LOQ level of 0.5 ng/ml could be established for cortisol. For corticosterone, linearity was evaluated in the range of 25–250 ng/ml, with 6 calibrator levels being included, i.e. 25, 50, 100, 150, 200, and 250 ng/ml. For evaluation of accuracy and precision, a low and high concentration level at resp. 25 and 200 ng/ml were involved, each on n= 6 replicates. The lowest level evaluated for accuracy and precision, i.e. 25 ng/ml, was established as the LOQ of the analysis method for corticosterone.

Results of the method validation experiments performed on chicken whole blood and mouse whole blood were all fully compliant with the acceptance criteria as given in the EC (2002/657/EC) and VICH (VICH GL49) guidelines.

Quantification of sheep whole blood samples was performed using the obtained mouse whole blood calibration curve of both corticosterone and cortisol.

**Table S2:**
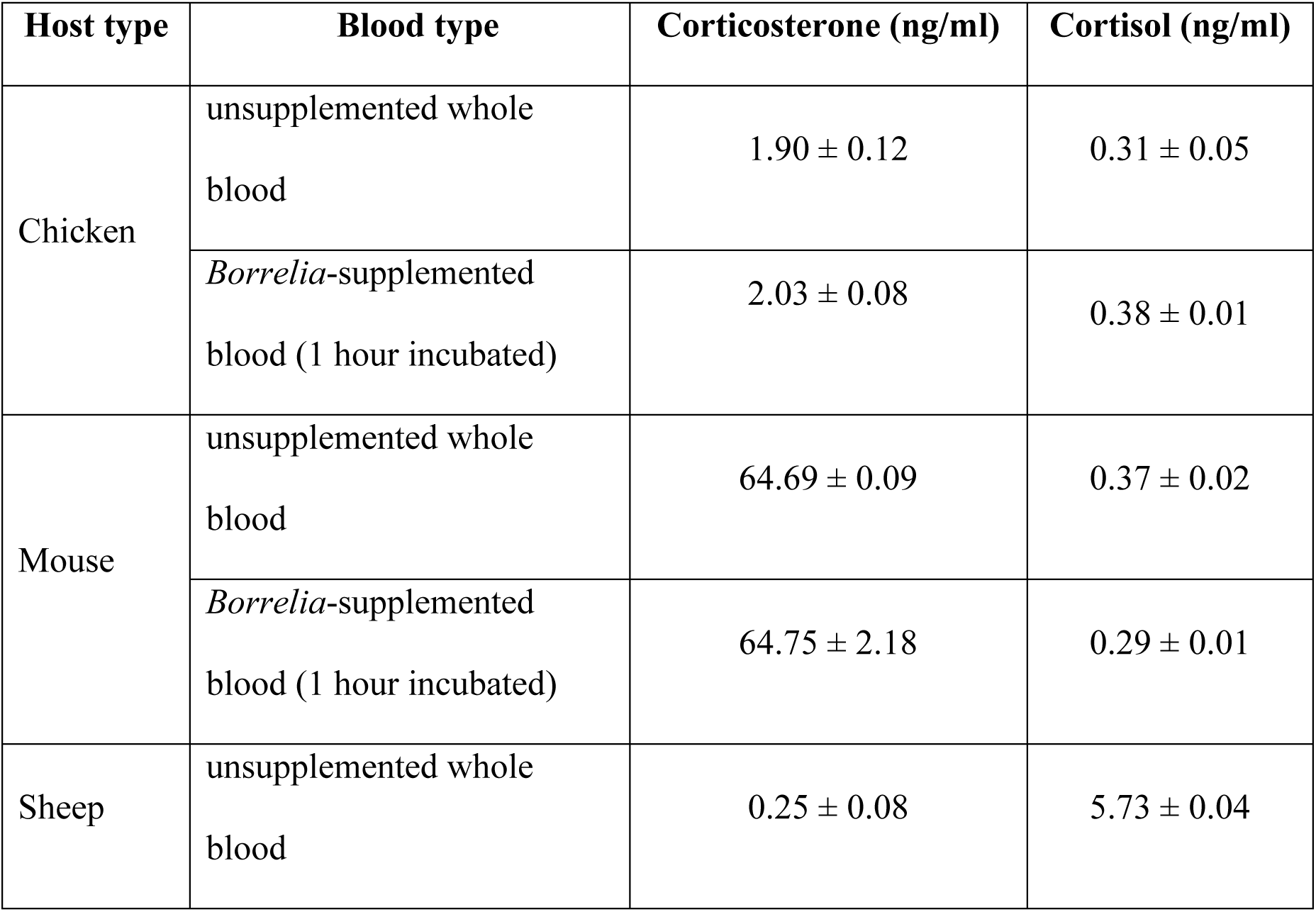
UPLC-MS/MS results of corticosterone and cortisol concentrations for unsupplemented whole blood and 1-hour incubated Borrelia-supplemented blood in chicken and mouse and unsupplemented whole blood in sheep. Mean ± SD is given, N=2.3.

The baseline glucocorticoid levels in mouse and chicken blood fall within the physiological range typically observed in laboratory mice and broiler chicken under non-stressed conditions (Lee et al., 2011; Gong et al., 2015; Hillebrecht et al., 2024; Scanes, 2016; Taban et al., 2016). These baseline values thus provide a meaningful reference point to evaluate the effects of glucocorticoid spiked blood.

There was no significant difference in glucocorticoid concentrations between unsupplemented bird blood and 1-hour incubated *Borrelia*-supplemented bird blood (corticosterone: t(1) = -1, p = 0.50; cortisol: t(1) = -2, p = 0.30). Similarly, no significant differences were observed in mouse blood (corticosterone: t(1) = -0.03, p = 0.98; cortisol: t(1) = 4, p = 0.16).

**Figure S1:**
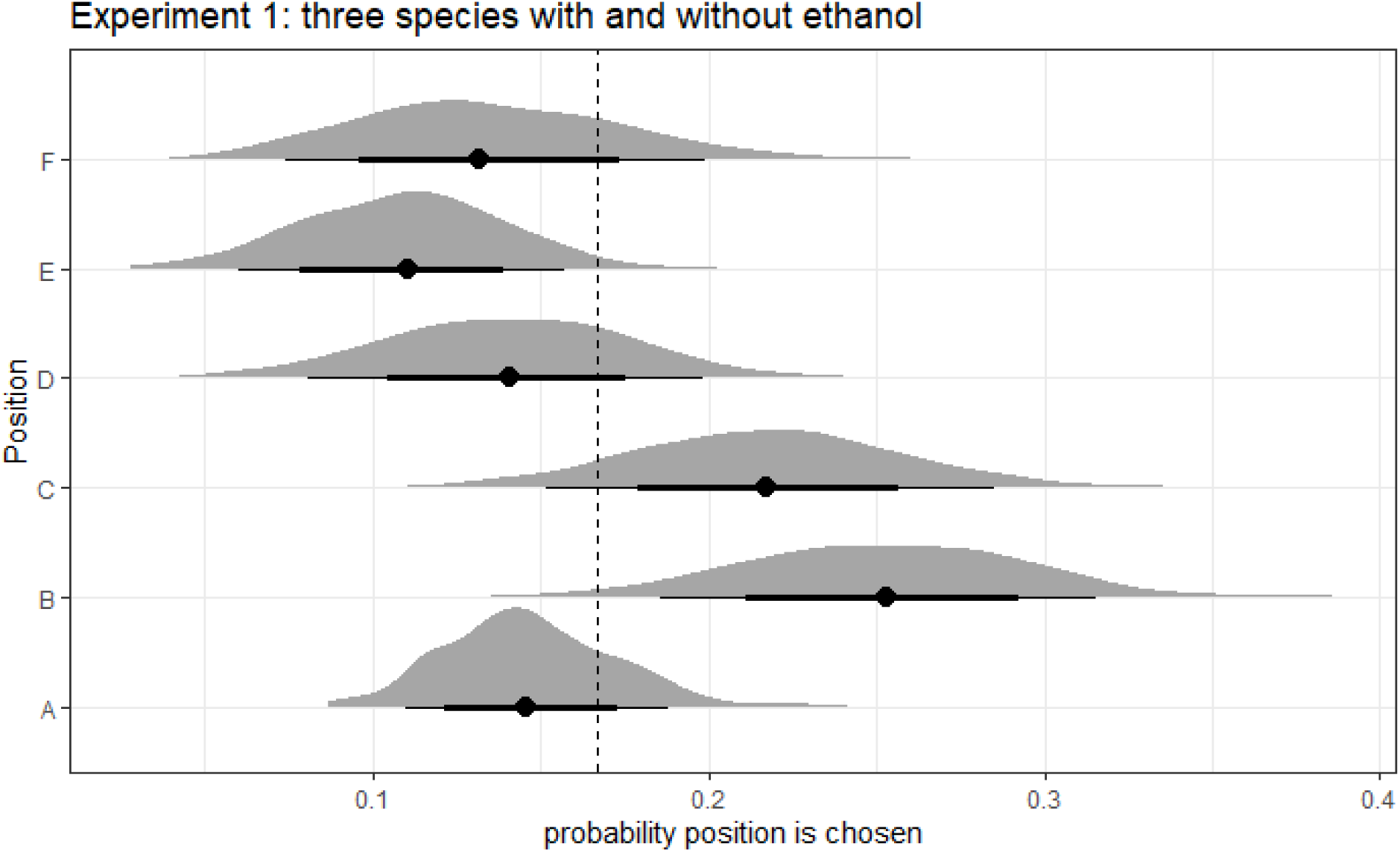
Tick preference for a position on the petri dish. Graphs show posterior distributions of estimated probabilities of a drop being chosen. The dot represents the median of the distribution with 90% (thin lines) and 68% (thick lines) credible intervals (CI). Dashed line is the expected probability when there is no preference.

**Figure S2:**
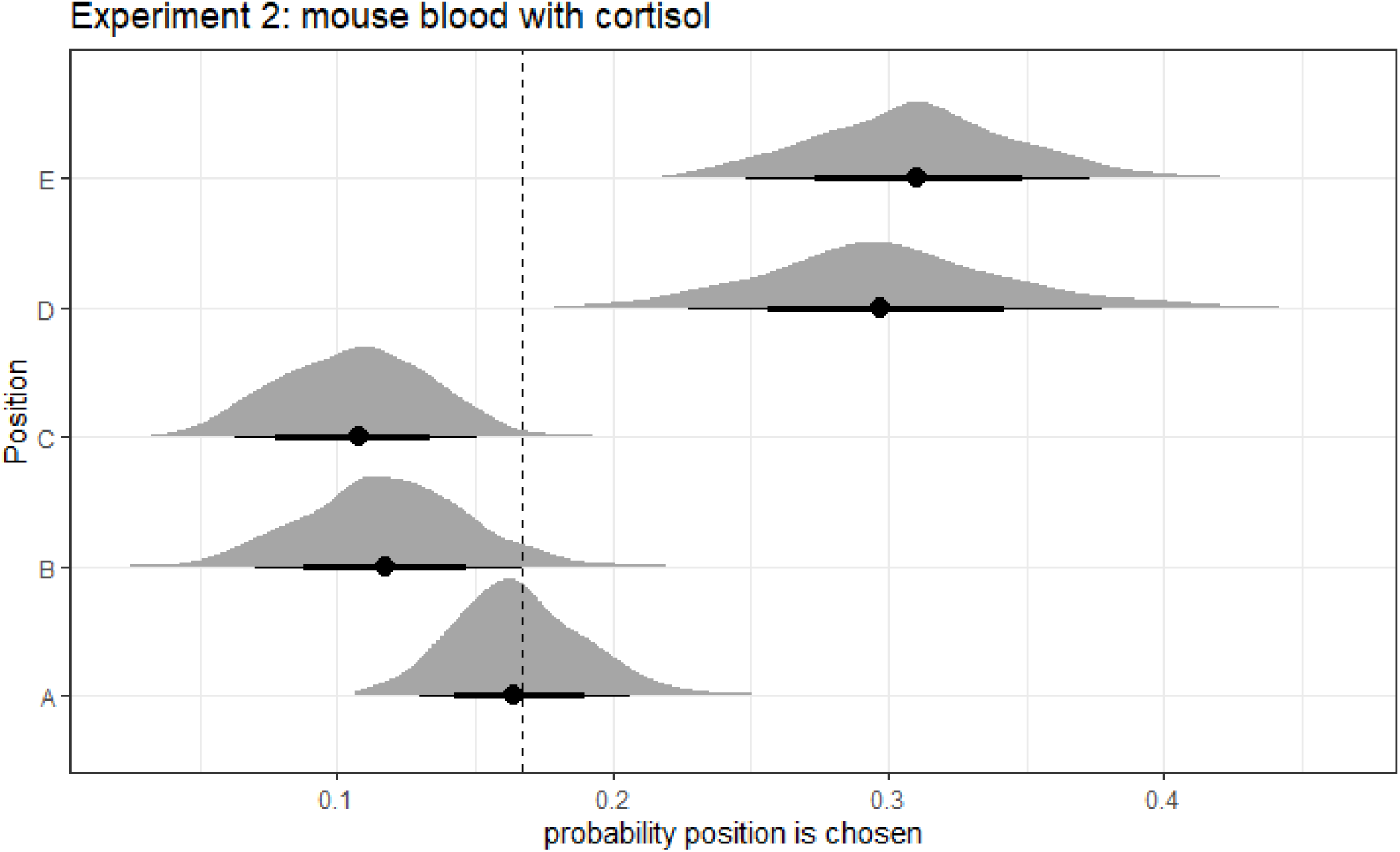
Tick preference for a position on the petri dish. Graphs show posterior distributions of estimated probabilities of a drop being chosen. The dot represents the median of the distribution with 90% (thin lines) and 68% (thick lines) credible intervals (CI). Dashed line is the expected probability when there is no preference.

**Figure S3:**
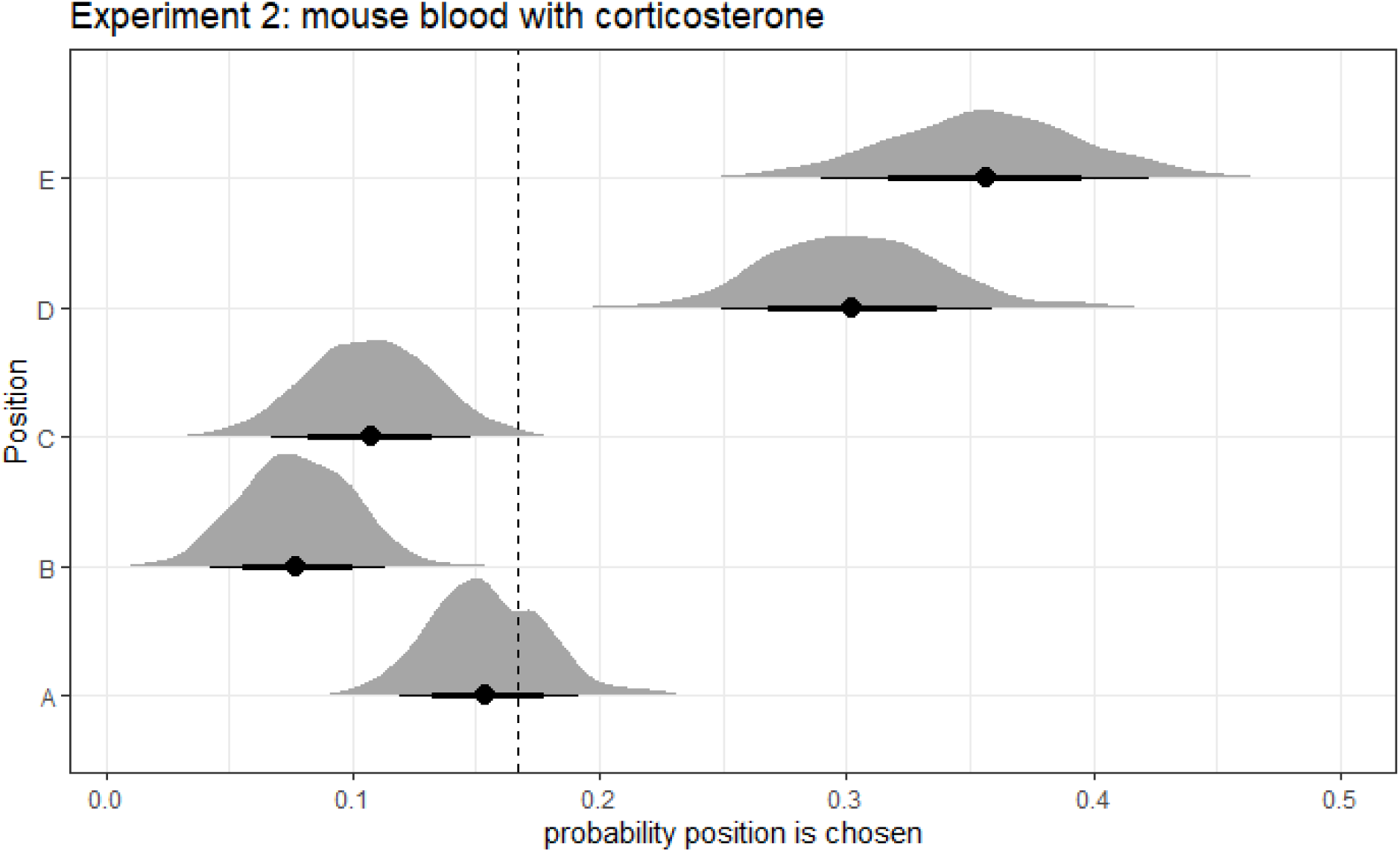
Tick preference for a position on the petri dish. Graphs show posterior distributions of estimated probabilities of a drop being chosen. The dot represents the median of the distribution with 90% (thin lines) and 68% (thick lines) credible intervals (CI). Dashed line is the expected probability when there is no preference.

**Figure S4:**
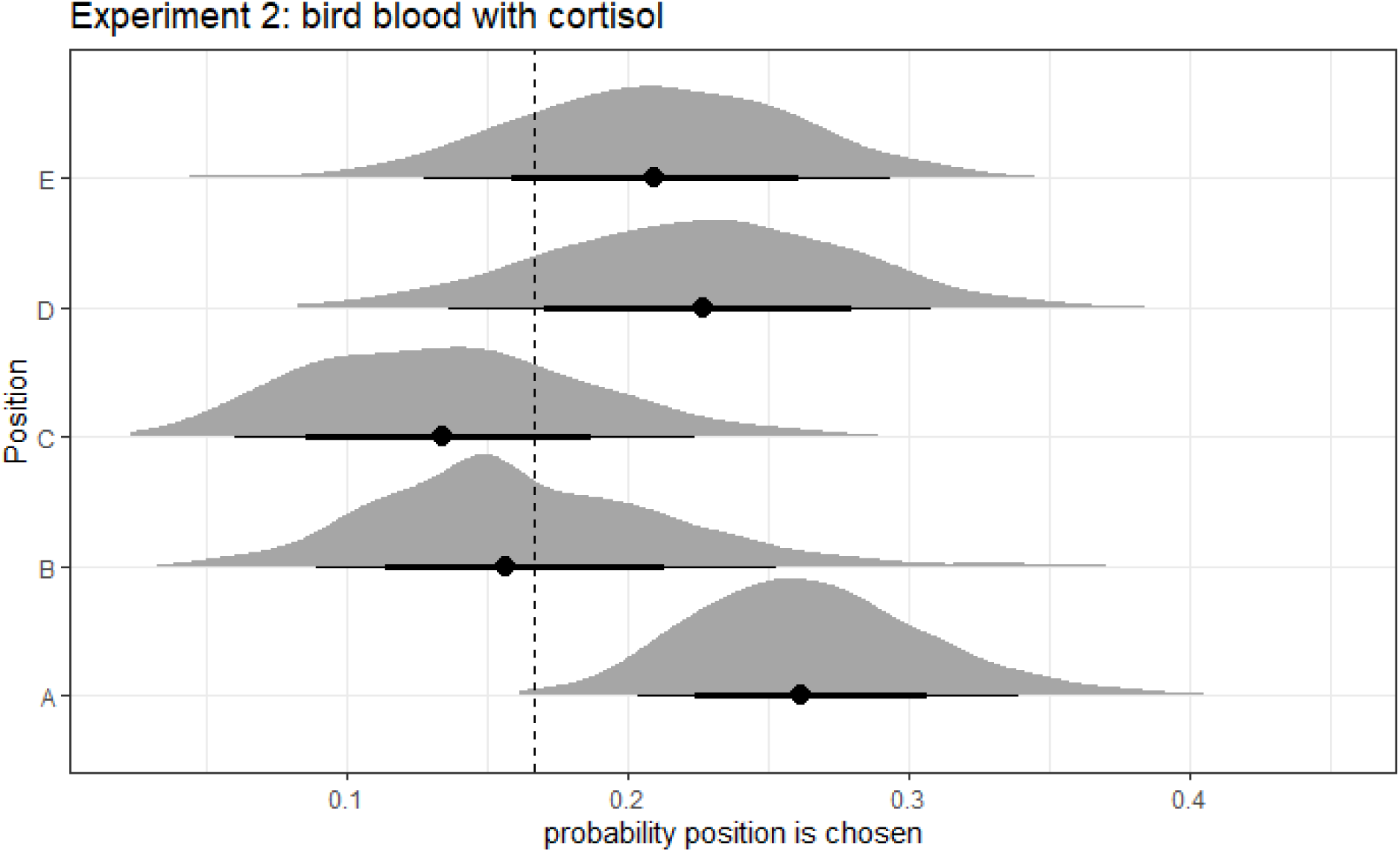
Tick preference for a position on the petri dish. Graphs show posterior distributions of estimated probabilities of a drop being chosen. The dot represents the median of the distribution with 90% (thin lines) and 68% (thick lines) credible intervals (CI). Dashed line is the expected probability when there is no preference.

**Figure S5:**
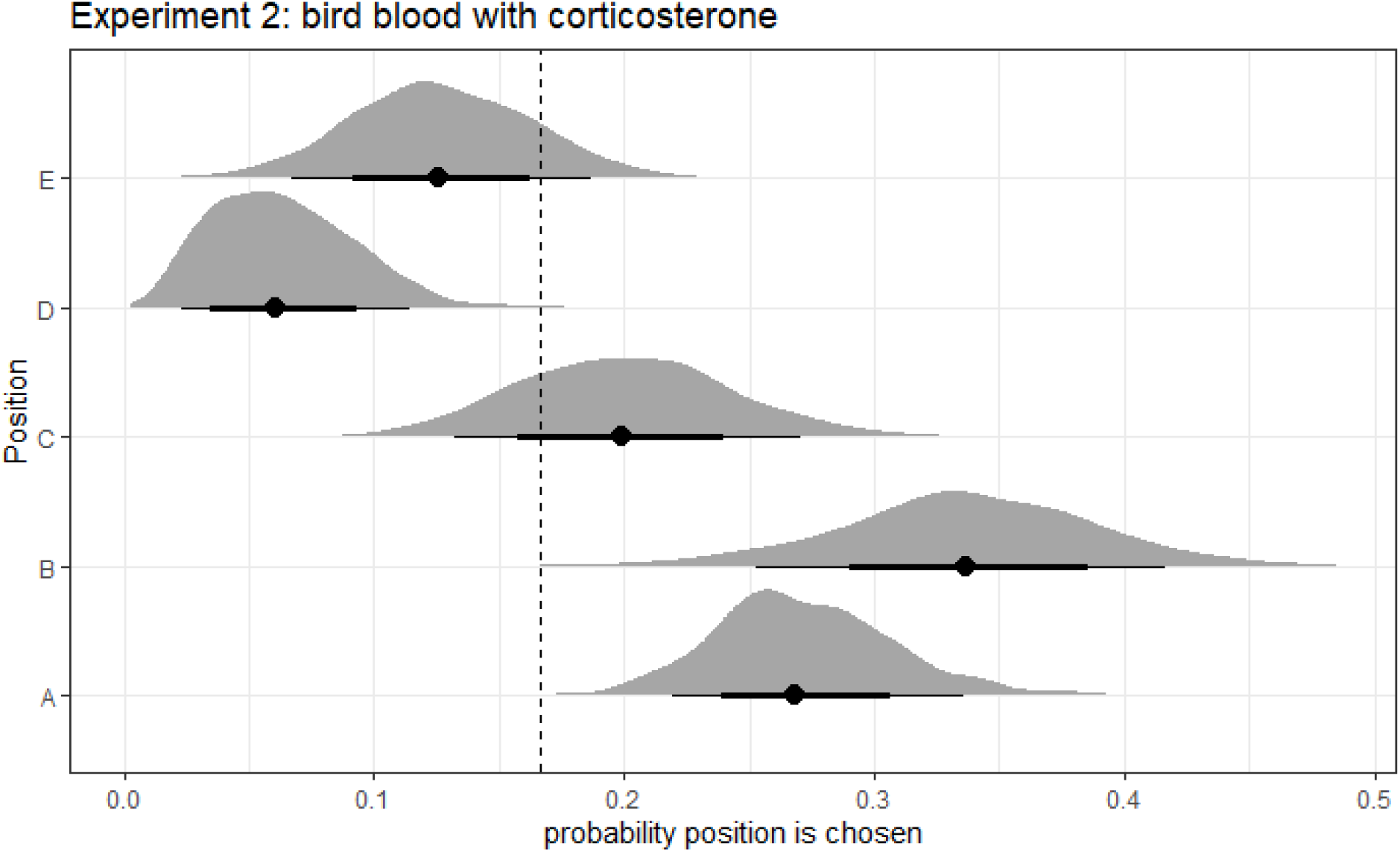
Tick preference for a position on the petri dish. Graphs show posterior distributions of estimated probabilities of a drop being chosen. The dot represents the median of the distribution with 90% (thin lines) and 68% (thick lines) credible intervals (CI). Dashed line is the expected probability when there is no preference.

**Figure S6:**
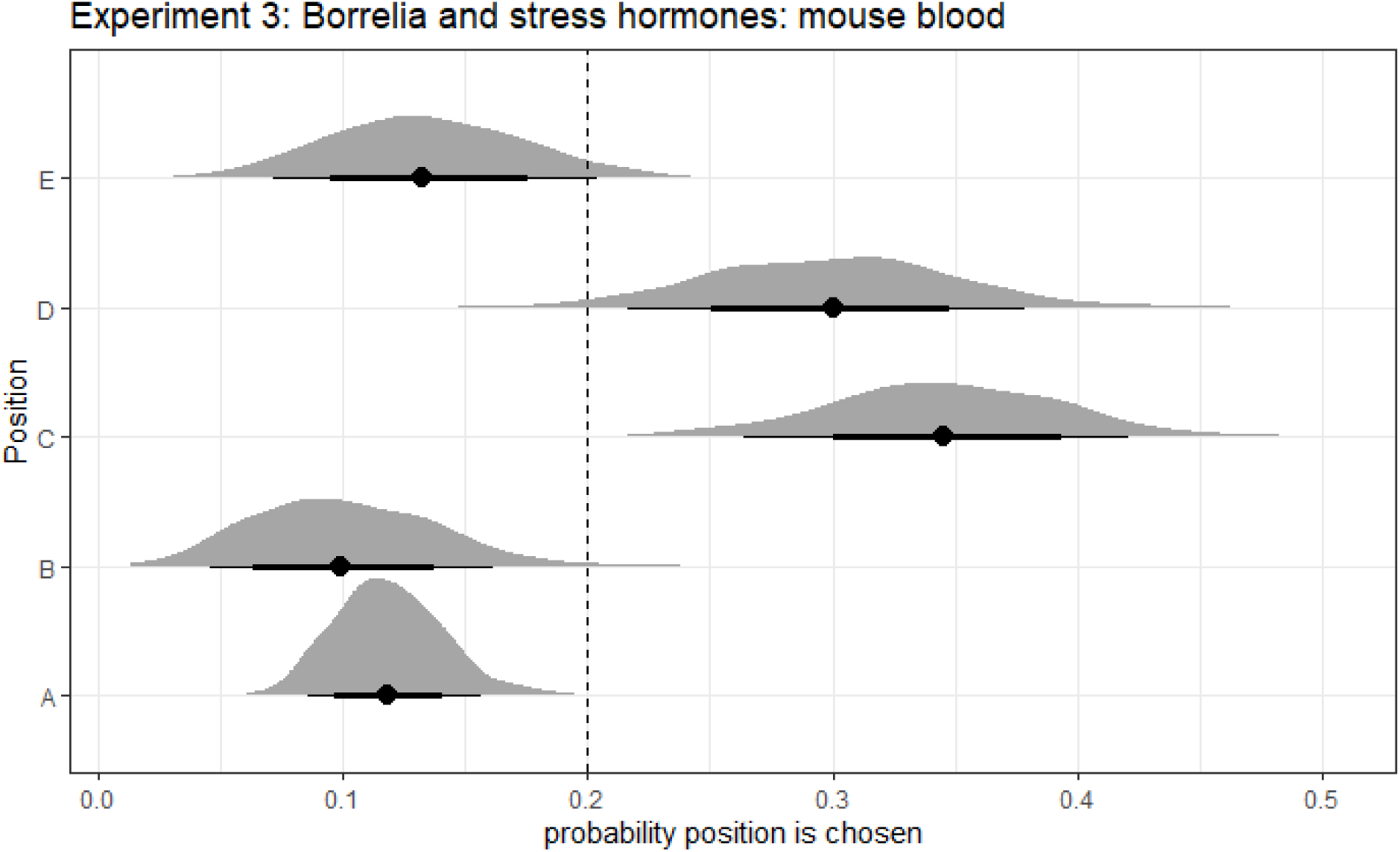
Tick preference for a position on the petri dish. Graphs show posterior distributions of estimated probabilities of a drop being chosen. The dot represents the median of the distribution with 90% (thin lines) and 68% (thick lines) credible intervals (CI). Dashed line is the expected probability when there is no preference.

**Figure S7:**
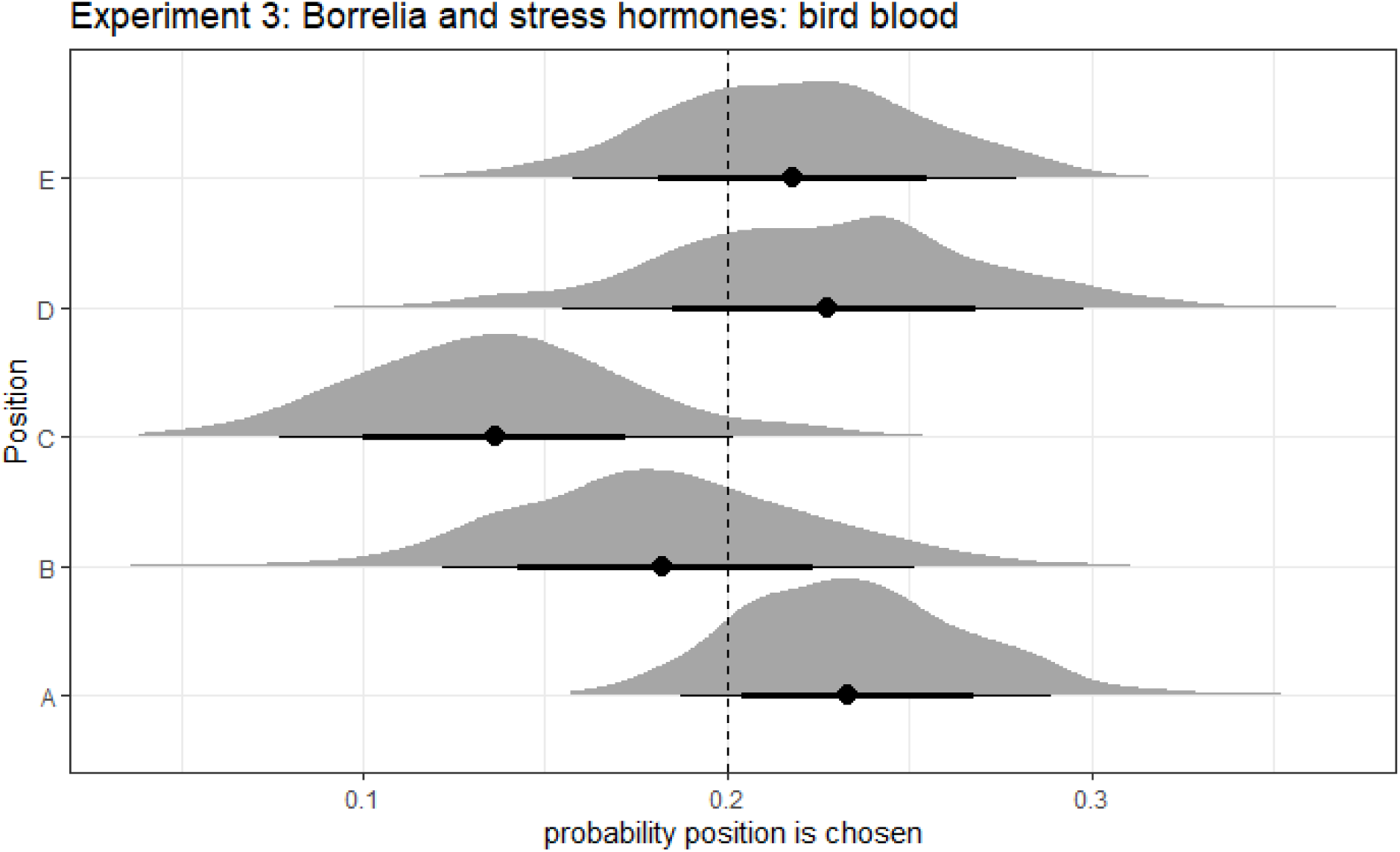
Tick preference for a position on the petri dish. Graphs show posterior distributions of estimated probabilities of a drop being chosen. The dot represents the median of the distribution with 90% (thin lines) and 68% (thick lines) credible intervals (CI). Dashed line is the expected probability when there is no preference.

**Table S3:**
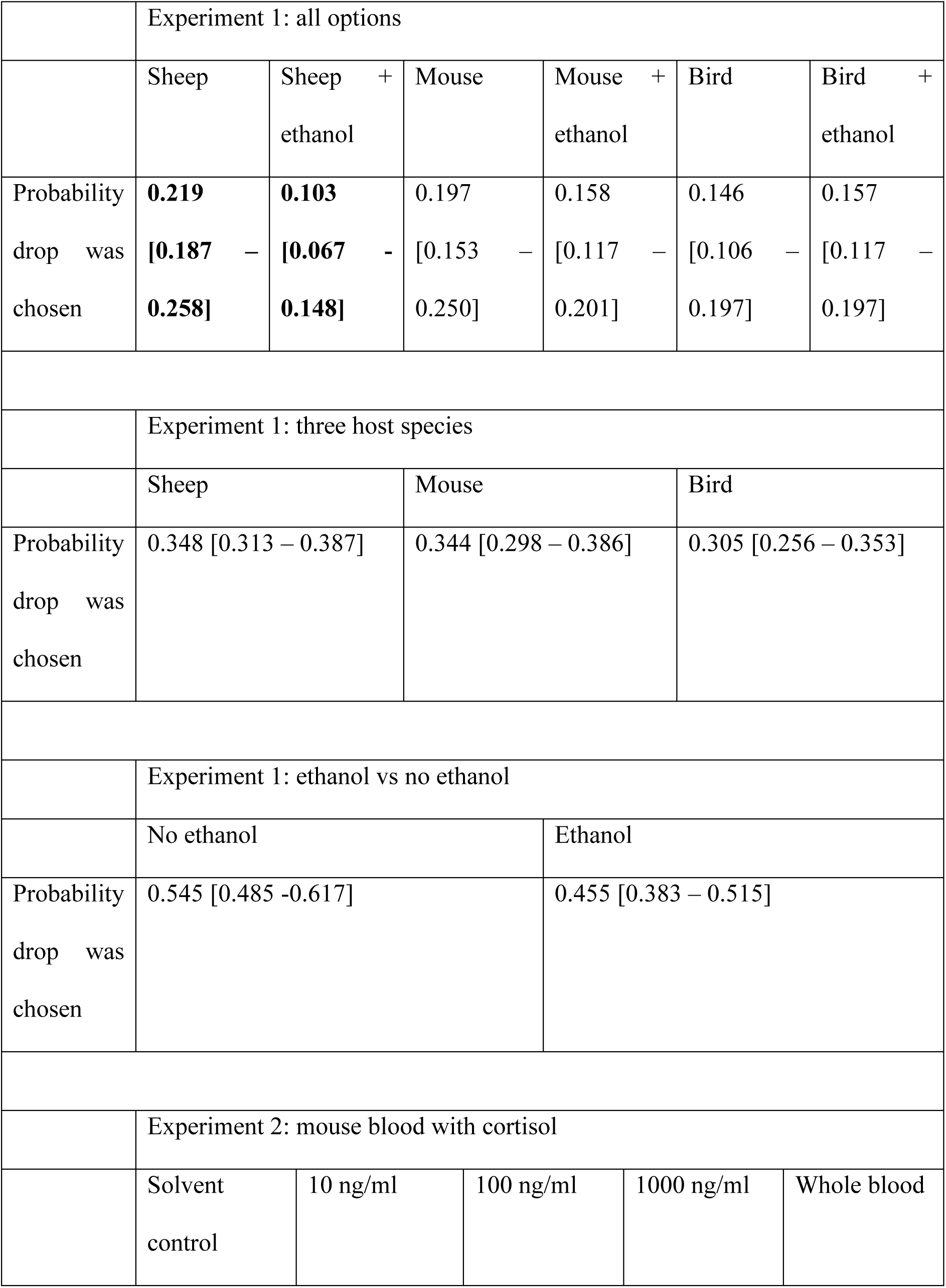
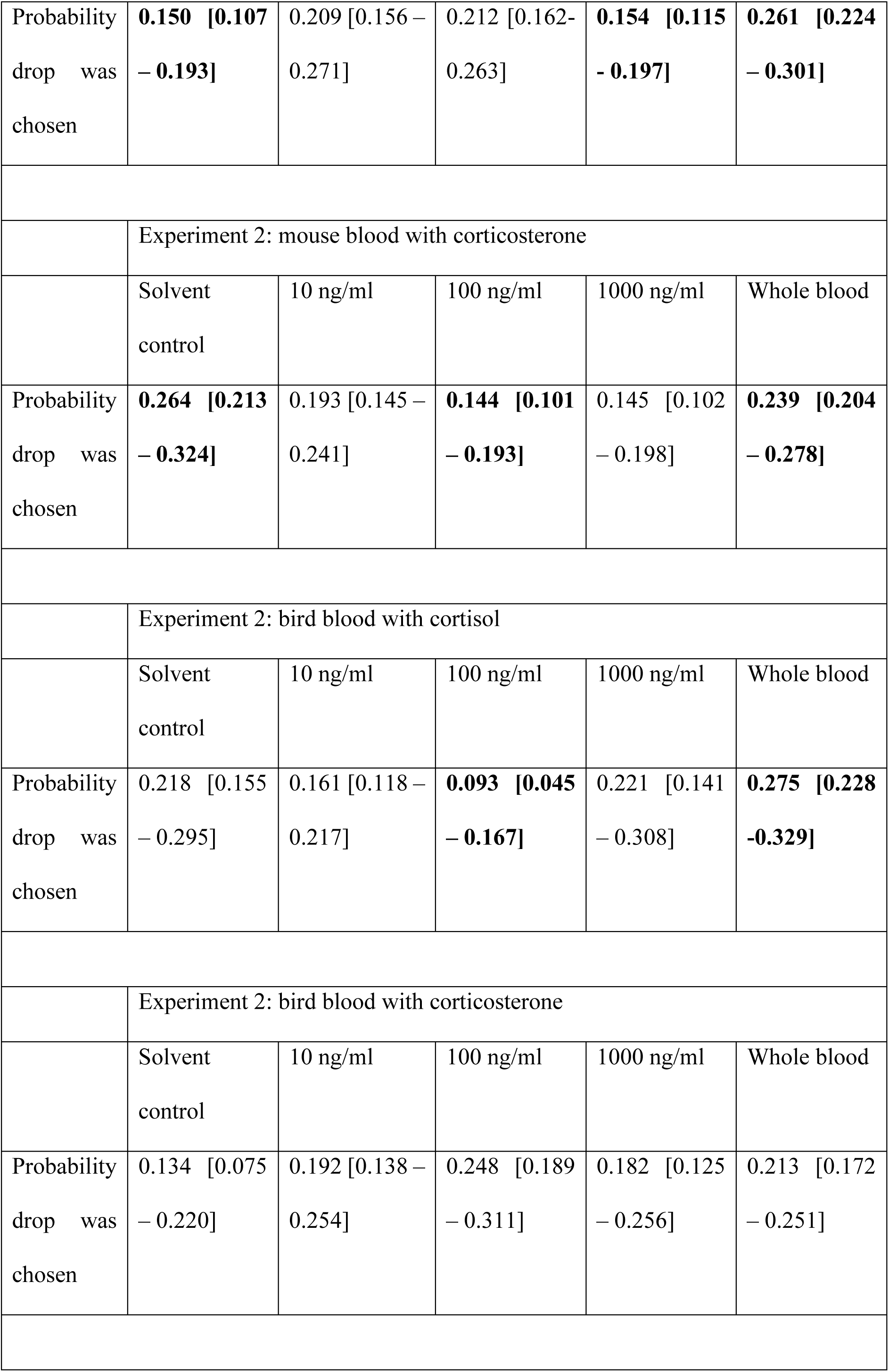
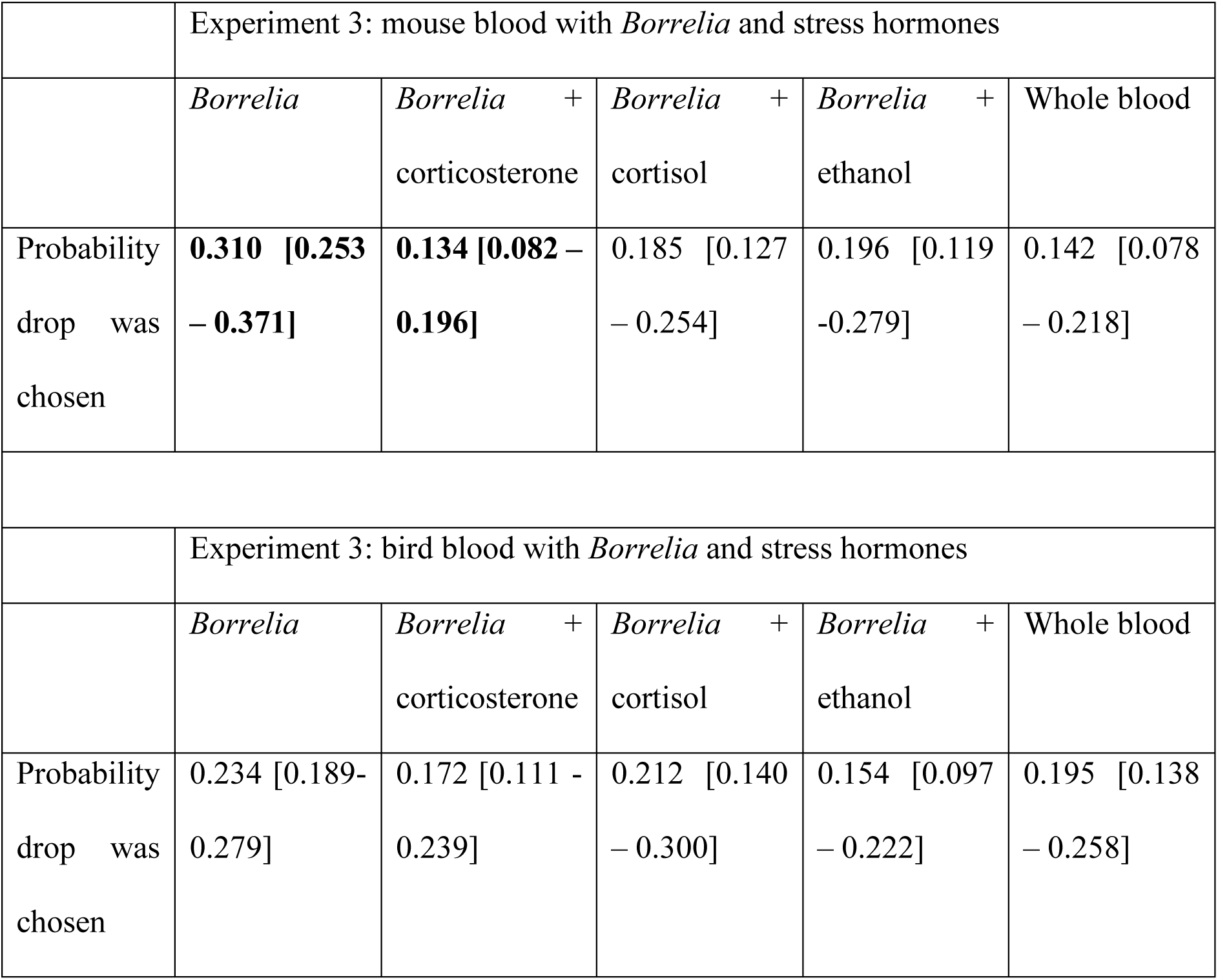
Median (+/- 68% quantile) expected probability a drop was chosen. In bold the ones that differ from the expected mean.

